# Adaptive LDA classifier enhances real-time control of an EEG Brain-computer interface for imagined-speech decoding

**DOI:** 10.1101/2023.12.12.571206

**Authors:** Shizhe Wu, Kinkini Bhadra, Anne-Lise Giraud, Silvia Marchesotti

## Abstract

Brain-Computer Interfaces (BCI) aim to establish a pathway between the brain and an external device without the involvement of the motor system, relying exclusively on neural signals. Such systems have the potential to provide a means of communication for patients who have lost the ability to speak due to a neurological disorder. Traditional methodologies for decoding imagined speech directly from brain signals often deploy static classifiers, that is, decoders that are computed once at the beginning of the experiment and that remain unchanged throughout the BCI use. However, this approach might be inadequate in effectively handling the non-stationary nature of electroencephalography (EEG) signals and the learning that accompanies BCI use as parameters are expected to change, all the more in a real-time setting. To address this limitation, we have developed an adaptive classifier that updates its parameters based on the incoming data in real time. We first identified optimal parameters (the update coefficient, UC) to be used in an adaptive Linear Discriminant Analysis (LDA) classifier, using a previously recorded EEG dataset, acquired while healthy participants controlled a binary BCI based on imagined syllable decoding. We subsequently tested the effectiveness of this optimization in a real-time BCI-control setting. Twenty healthy participants performed two BCI-control sessions based on the imagery of two syllables, using a static LDA and the other the adaptive LDA classifier, in randomized order. In this real-time BCI-control task, the adaptive classifier led to better performances than the static one. Furthermore, the optimal parameters for the adaptive classifier were closely aligned in both datasets acquired with the same syllable imagery task. These findings highlight the effectiveness and re-liability of adaptive LDA classifiers for real-time imagined speech decoding, and its interest for non-invasive EEG-based BCI notably characterized by low decoding accuracies.

## 1. Introduction

Speech stands as a distinct human capability, pivotal for both communication and personal expression. Neurological disorders such as locked-in syndrome (LIS) and aphasia *(1)* can impair the ability to communicate with the external world, while leaving cognitive functions, including consciousness, intact *(2)*. Harnessing these residual cognitive capabilities has the potential to restore communication, thereby enhancing patients’ overall quality of life. A promising approach is to use a speech brain-computer interface (BCI), a system able to provide the user with a means of communication by decoding neural activity related to speech intentions directly from brain recordings, without the involvement of the motor system. Recently, the field has witnessed major advances in decoding from electrodes implanted intracranially *(3–6)*. Previous studies have demonstrated the feasibility of synthesizing attempted speech from cortical signals in offline analyses *(7–10)*, and recent breakthroughs have shown impressive performance in real-time decoding *(11–14)*, reaching a decoding rate of 78 words per minute *(14)*.

Despite its effectiveness, decoding attempted speech is precluded in patients with language disorders in which the damage is located upstream with respect to the motor representations, such as in aphasia. In these patients, a more appropriate approach would be to decode imagined (i.e. covert) rather than attempted speech. Speech imagery is defined as the silent, internal generation of speech elements without any audible output or associated physical articulation *(3)*. Speech imagery entails less cortical activation and different cortical representations than overt, articulated speech *(15–19)*. These characteristics make covert speech more difficult to decode from neural signals, especially in real-time, and suggest that methodologies optimized for overt speech-based BCIs cannot seamlessly be translated to systems designed for speech imagery. Thus, it is not surprising that BCI based on covert rather than overt speech lags behind in terms of research breakthroughs and developments. Real-time speech synthesis with acceptable intelligibility from speech imagery remains an important goal to be achieved *(5)*.

Despite these challenges, previous studies have delved into the complexities of covert speech decoding, using both intracranial recordings *(6,20,21)*, and non-invasive techniques such as EEG *(22–24)*. These studies involved the imagery of different types of speech units such as vowels *(25,26)*, syllables *(27,28)*, words *(21,29–31)* and sentences *(32)*. Only two previous BCI studies based on surface EEG recordings have so far attempted to decode imagined speech in real-time and have shown the limited effectiveness of current decoding approaches *(23,33)*. Many more studies have targeted the decoding of imagined speech with offline analyses with the main goal of enhancing current classifiers, but none have tested their effectiveness in improving BCI-control in real time. Yet, this is a crucial aspect to consider due to EEG non-stationarity *(34)* that can hamper decoding in real-time. Significant variations in the signals occur throughout the course of an experiment due to different reasons. These include the hardware employed to record brain signals through EEG (e.g. drying of the conductive gel or shifts in the position of the electrodes), physiological fluctuations in cognitive states (such as attention, motivation, and vigilance levels, *(35)* and neural changes due to short-term brain plasticity resulting from practicing the task. In addition, BCI experiments can be affected by yet another source of variability, namely the experimental discrepancy between the *offline training* part in which data are acquired to calibrate the classifier without providing feedback to the user (open-loop), and the *online control* part in which the user is given a feedback in real-time (closed-loop). For all these reasons, a classifier developed at the beginning of the experiment based on offline data might not be able to optimally decode EEG signals in real-time.

To mitigate the effects of non-stationarities, the use of adaptive machine learning techniques in BCI systems emerges as a promising solution. Compared to static classifiers that remain constant throughout the entire duration of the experiment, adaptive BCI systems continuously update either their decoding features or classifier parameters based on new incoming signals during real-time control *(36,37)*. Unlike their static counterparts, these approaches enable a co-adaptive calibration between the user and the machine during online control, enhancing both BCI controllability *(38,39)* and signal decidability. The effectiveness of these decoding methods has been established in previous motor-imagery BCI studies *(39–41)*, which highlighted that decoding improvements with an adaptive classifier does not only help users with poor BCI-control but can also speed up the learning curve in good BCI users *(38,39,42)*.

In terms of feature extraction, methodologies such as adaptive common spatial patterns *(43,44)*, supervised feature extraction *(45)*, adaptive autoregressive parameters *(46)* and adaptive wavelet packet transform *(47)* have shown their effectiveness in improving the discriminability of decoding features. The other approach consists in using adaptive classifiers that update their parameters (e.g. the weights assigned to each feature in the linear discriminant hyperplane) whenever new data is available. These methods have proven to consistently perform well with non-stationary EEG signals *(36,48)*. Different kinds of linear classifiers can be translated into their adaptive counterparts, including linear discriminant analysis (LDA) *(35,49,50)*, support vector machines *(51)* and Bayesian classifiers *(52)*.

Given the success of adaptive classifiers in motor imagery BCIs, similar benefits can be expected in speech imagery BCI *(22)*, where decoding is notably difficult. We thus conducted a BCI-control study to directly assess the advantage of using an adaptive rather than a static LDA classifier to decode the imagery of two syllables from EEG signals. We first fine-tuned the parameters of adaptive LDA classifier with open-loop simulations performed using a pre-recorded dataset. Subsequently, we tested for performance improvement in real-time by comparing BCI-control using the static and the adaptive LDA with these optimized parameters in a group of twenty healthy volunteers, each of whom underwent two BCI control sessions, using each classifier on separate days. Last, we explored an alternative approach for the optimization of adaptive classifiers based on individualized tuning of the classifier parameters.

## 2. Materials and Methods

### 2.1 Participants

We first used EEG data from a previous study (Study 1) conducted in our laboratory *(33)*, comprising 15 healthy volunteers (5 women; average age: 23.9 years, SD ± 2.3, age range: 19-29 years) to conduct decoding simulations described in Section 2.5 “Simulation analysis”. In a second step (Study 2), we recruited twenty healthy participants (13 women; average age: 25.2 years, SD ±3.1, age range: 20-30 years). The two groups included different individuals. Both studies were approved by the local Ethics Committee (Commission cantonale d’éthique de la recherche, project 2022-00451) and were performed in accordance with the Declaration of Helsinki. All participants provided written informed consent and received financial compensation for their participation.

### 2.2 Experimental paradigm

All participants of Study 2 underwent two BCI-control sessions on different days, spaced at least by 3 weeks to avoid potential learning effects, that could represent a confounding factor when evaluating differences in performance between the two sessions. During these sessions, either a static or an adaptive LDA classifier was used in counterbalanced order across participants. BCI-control sessions were conducted in a room shielded optically, acoustically, and electrically, each of which lasted approximately 3 hours, amounting to a total of 6 hours of experimental time per participant. Participants were orally briefed about the overall procedure (see section 2.2.1 and section 2.3) prior to the experiment.

#### 2.2.1 Syllable imagery

As in Study 1, the goal of Study 2 was to decode the imagery of two syllables /fɔ/ and/gi/ (presented in written form on the screen during the experiment as “fo” and “ghi”). They were selected based on their distinct phonological and acoustical attributes with respect to the consonant manner (fricative vs plosive), place of articulation (labiodental vs velar), vowel place (mid back vs high front) and rounding (rounded vs unrounded). Previous research indicated that distinct phonetic features elicit different neural responses *(18,53–55)*. We assumed this choice would maximize the discriminability of the neural signals elicited by the imagery of the two syllables, hence facilitating decoding.

Participants were instructed to engage in the imagery of one of the two syllables, concentrating on the kinesthetic sensations associated with actual pronunciation. They were instructed to avoid auditory or visual representations, such as internal hearing or visualizing the syllable in written form. Clear instructions were provided to refrain from any physical movements during the imagery process, with particular emphasis on avoiding facial movements and any subvocalization. They were instructed to maintain focus during the syllable imagery and were allowed to take breaks if needed.

### 2.3 EEG acquisition and BCI loop

#### 2.3.1 EEG recording

EEG data were acquired using a 64-channel ANT Neuro system (eego mylab, ANT Neuro, Hengelo, Netherlands) with a 512 Hz sampling rate. Electrode AFz served as the ground, and CPz was used as the reference. Impedances for the ground and reference electrodes were consistently maintained below 20 kΩ, while the impedances for all other electrodes were kept under 40 kΩ throughout the experiment.

Neural signals were collected through the Lab Streaming Layer system (LSL, https://github.com/sccn/labstreaminglayer).

While the EEG was being recorded and the BCI was in operation, participants remained seated comfortably in front of a computer screen with their hands resting on their lap.

#### 2.3.2 BCI loop

The BCI loop was developed upon the *NeuroDecode* framework (Fondation Campus Biotech Geneva, https://github.com/fcbg-hnp/NeuroDecode.git) already employed in previous BCI experiments *(33,56,57)*.

The BCI experiment included three phases: (a) *offline training*, (b) *classifier calibration*, and (c) *online control*.

During the *offline training*, participants performed the syllable imagery without receiving real-time feedback while data was being recorded. The offline session comprised 4 blocks of 20 trials in Study 1, and 3 blocks of 30 trials each in Study 2, and lasted for 25-30 minutes. Within each block, both syllables were represented an equal number of times in randomized order. In both studies, each offline trial began with a text indicating the trial number (1 s), followed by a fixation cross (2 s), after which a written prompt indicated which of the two syllables the participants had to imagine pronouncing (2 s). After the cue disappeared, an empty battery appeared on the screen and immediately started to fill up until fully filled across the subsequent 5 seconds. Participants were instructed to start imagining pronouncing the cued syllable as soon as the battery appeared and to stop once the battery was full. They were instructed to keep a consistent pace while imagining repeating the syllable throughout the entire experiment. Subsequently, participants were given a rest period, while the word “Rest” (4 s) was displayed on the screen.

During the *classifier calibration* phase, offline data were used to train an LDA classifier. A standard LDA classifier uses a hyperplane to separate data from different classes by finding the optimal projection that maximizes between-classes distance while minimizing within-classes distance *(58)*. The choice of this classifier was motivated by several studies showing the effectiveness of the adaptive version in improving BCI performance as compared to the static version *(35,39,49,50,59,60)*.

The LDA classifier was designed to distinguish between the two syllables based on features derived from the EEG power spectral density (PSD) in the 1-70 Hz frequency range, with a 2 Hz interval, and from all EEG electrodes except the reference channel and the two mastoid electrodes (leaving 61 channels). The PSD was computed using a 500 ms sliding window with a 20 ms overlap. The classifier’s performance was determined through an 8-fold cross-validation accuracy (“CV accuracy”).

During the *online control* phase, participants imagined pronouncing either one or the other syllable as in the *offline training* phase, except that this time their brain activity was analyzed in real-time by the LDA classifier and the output of the decoder used to control the battery filling in real-time. Participants were informed that the visual feedback would be updated (around every 50 ms) according to the classifier’s probability output: if the output corresponded to the cued syllable the battery filling increased, otherwise the battery filling decreased. The battery filling real-time control continued until the battery was either fully charged or after a 5-second timeout. Participants were instructed to use the same syllable-imagery strategy as during the offline training.

During the “*static classifier”* session, the parameters obtained in the calibration phase were used throughout the entire online control phase (see Figure 2a). In contrast, during the “*adaptive classifier”* session, the LDA classifier was updated iteratively using features derived from newly acquired data (Figure 2b), every time a new trial from both classes (i.e. the two syllables) becomes available.

**Figure 1:**
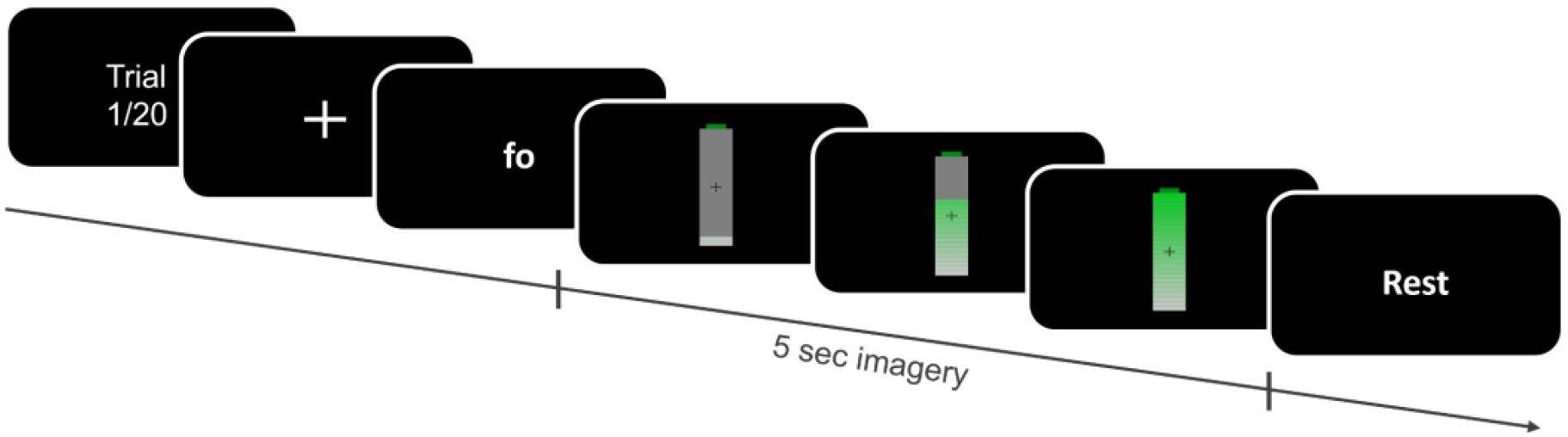
Experimental Procedure Overview (Studies 1&2). Each trial began with the display of the trial number, followed by a fixation cross. Participants then received a textual cue indicating the syllable to imagine (fo or ghi). During the *offline training* phase, the imagery task lasted for 5 seconds, while in the *online control* phase, it continued until the battery icon on the screen was filled or until a 5 second timeout. Participants were given 4 seconds of rest at the end of each trial.

**Figure 2:**
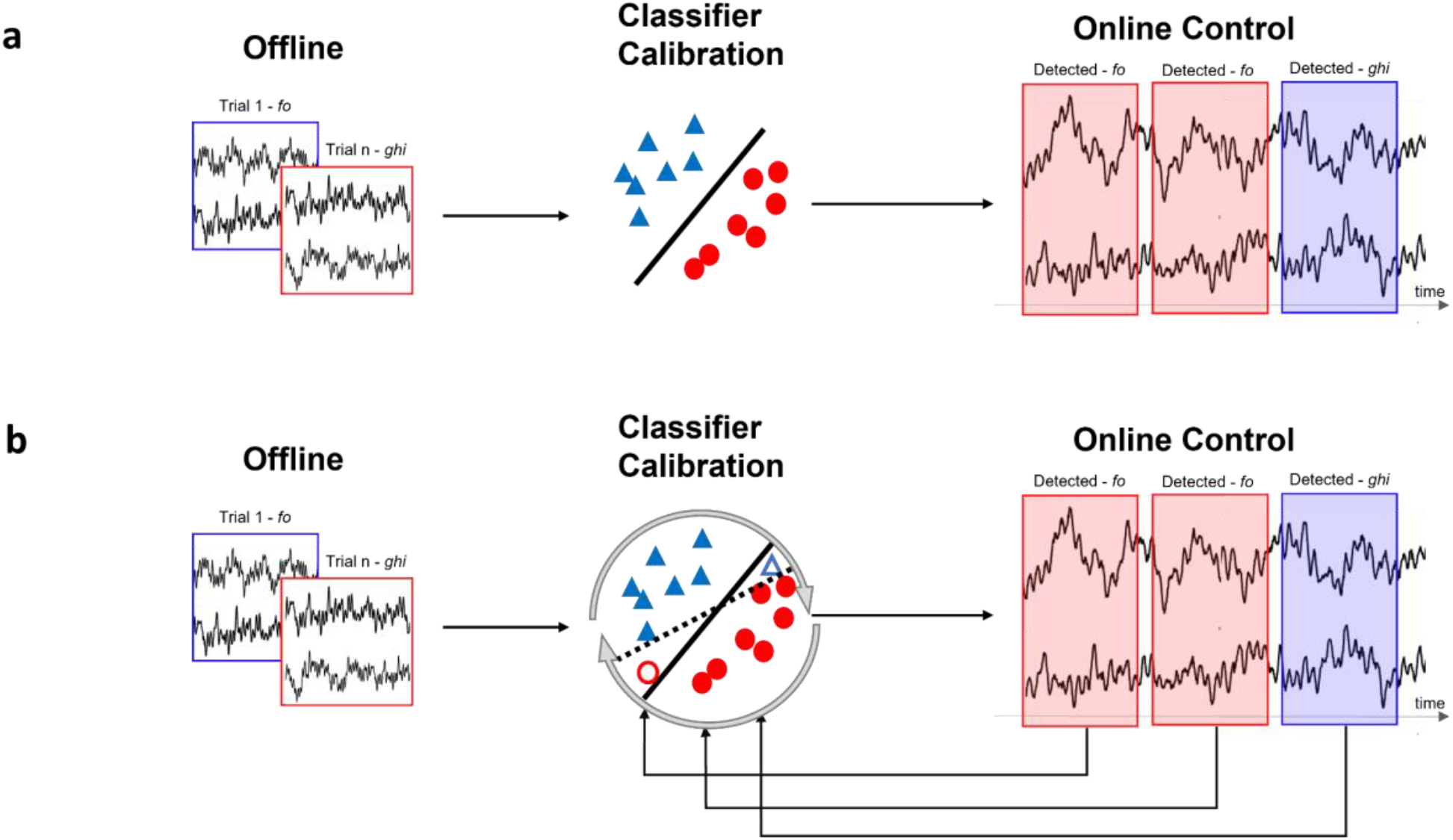
Schematic representation of the static and adaptive classifier. (a) Static: the LDA classifier was built based on the dataset acquired during the *offline training* phase and remained unchanged throughout the entire *online control* phase. (b) Adaptive: an initial classifier was built as in (a), and subsequently updated iteratively during the *online control* phase using features derived from new incoming data in real-time. Red circles and blue triangles indicate features belonging to the two classes; empty shapes denote features from newly acquired data (during the *online control* phase) that are used to update the classifier. The solid lines in the Classifier Calibration step represent the separating hyperplane obtained with the initial static classifier, while the dashed line illustrates the updated classifier after the new data are integrated.

### 2.4 Adaptive LDA classifier

For a binary classification, the LDA operates on the principle of distinguishing between two classes by assuming that both follow a normal distribution. This assumption is applied to the offline dataset by retrieving the mean vectors and covariance matrices for each of the two classes. Furthermore, it is assumed that both classes are characterized by a common covariance matrix *(58)*. The LDA classifier can ascertain the data label using Equations (1) to (3). The value *D(x)* represents the differential distance of the feature vector *x* from the separating hyperplane, characterized by its normal vector *w* and bias *b*. The parameters *w* and *b* can be expressed in terms of the inverse covariance matrix ∑^-1^ (2) and the mean of each class *μ*_*1*_ and *μ*_*2*_ (3). In our case, the feature vector *x* is derived from one sliding window of neural data relative to the imagery of one syllable. The sign of D(x) determines the class (i.e. the syllable imagined) to which the observed feature vector *x* is assigned.

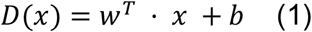

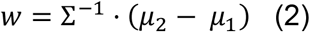

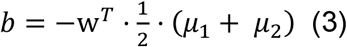

One way to achieve adaptive classification consists in updating the parameters *w* and *b* based on newly acquired data in real-time, during the *online control*. To do so, we employ an algorithm that consists in changing at each time *t* (in our case when each class has a new trial) the value of the mean *μ* of each class *i* (equal to 1 or 2, for a two-class problem) and the common inverse covariance matrix ∑^-1^ according respectively to Equations (4) and (5) *(50)*. In these equations, an updated coefficient (UC) is used to set the respective contribution of the previous model’s values of the mean *μ*_*i*_(t-1) and covariance matrix *∑(t-1)*, as well as the current features *x(t)* and ultimately to update the current mean *μ*_*i*_*(t)* and covariance matrix *∑(t)*. The values of the UC (one for the mean UC_μ_ and the other for the covariance matrix UC_∑_), between 0 and 1, set the updating speed of the adaptive classifier. A value nearing zero indicates minimal influence of new samples on the model, whereas a value approaching one results in a stronger reliance on newly acquired samples as compared to old ones and leads to a faster update.

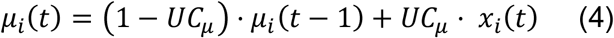

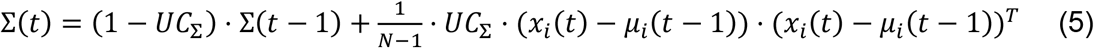

The inverse of the covariance matrix ∑(t)^-1^ was computed by applying the Woodbury matrix identity (equation (6) and (7), *(61)*:

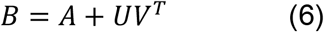

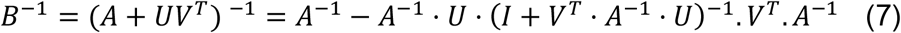

Using these equations and by integrating values from *x(t), ∑(t-1)*^*-1*^, *μ*_*x*_*(t-1)*, UC_∑_ and UC_μ_, the inverse of the covariance matrix *∑(t)*^*-1*^ and *μ*_*i*_*(t)* for each class were updated iteratively in real-time as follows (equations (8)-(11)):

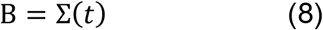

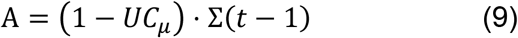

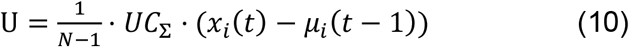

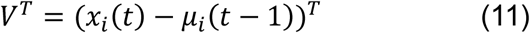

### 2.5 Simulation analyses to optimize the adaptive LDA classifier

While most parameters in the adaptive LDA are data-driven (e.g. *x, ∑*^*-1*^, *μ*_*x*_), UC_∑_ and UC_μ_ are constants that need to be pre-specified. To identify the optimal UC values, specific to the syllable imagery task, to be used in the adaptive LDA classifier, we conducted classification simulations on the Study 1 EEG dataset *(33)*. Importantly, the binary /fɔ/ vs. /gi/ syllable imagery task was exactly the same in Study 1 (optimization) and in Study 2 (static vs. adaptive strategy comparison). The Study 1 dataset consisted of EEG recordings from 15 healthy participants who trained to control a speech-BCI daily over five consecutive days, yielding a total of 75 datasets.

We probed a series of values for the UC_μ_ and the UC_∑_, ranging from 0.4 * 2^-14^ to 0.4 in increments of powers of two (15 values in total). The upper bound was set to prevent the update speed from being excessively rapid. To identify the best UC value range, we evaluated the predicted accuracy (PA) obtained through simulations of classification using every possible combination of UC_μ_ and UC_∑_ values (UC-pair), for each of these pre-recorded datasets. To compute the PA, an initial LDA classifier was constructed using data acquired during the *offline training* phase (80 trials). Then, for a given UC-pair, the classifier was updated with data from the *online control* for each new trial and class. To classify each trial, we split the trial into 500 ms sliding windows with a 50 ms overlap, simulating the real-time process during *online control*. The decoding accuracy for the cued syllable was computed for each sliding window and probabilities from all decoding accuracies were averaged. The trial was labeled as correctly or wrongly classified based on the average value being above or below a 50% chance level. Finally, the PA for that specific UC-pair was computed as the percentage of correct trials over the total number of trials. This led to a 15x15 matrix, “adaptive predicted accuracy” (adaptive PA) matrix, in which the x-axis represents the range of possible values for the UC_∑_ and the y-axis the range of UC_μ_ values, and each element of the matrix indicates the adaptive PA.

To assess the effectiveness of this approach we compared the adaptive PA with the PA obtained with a static LDA classifier. To compute the static PA, we apply an approach analog to the one described above: first, we computed a classifier based on the *offline training* dataset and subsequently used it to predict each trial in the *online control*, this time without updating the classifier parameters. We then considered the percentage of trials correctly classified to obtain the static PA for each of the 75 EEG datasets. Of note, differently from the adaptive PA matrix, the static classifier did not involve different UC values, thus there was only one static PA value per dataset.

The adaptive PA matrix and the static PA were used to derive three new matrices from which we selected the UC-pair: the Modulated PA, the Consistency Score, and the Modulated Bias matrices.

#### 2.5.1 “Modulated predicted accuracy” matrix

To compare the PAs obtained with the static and adaptive classifier and identify the optimal UC-pair, we derived, for each dataset, the Modulated PA matrix by subtracting the static PA from the corresponding a-PA matrix, separately for each of the 75 datasets. We then computed the average of all Modulated PA matrices. Finally, we identify the UC-pair (UC_μ_ and UC_∑_ values) that led to the highest PA increase with the adaptive as compared to the static classifier.

#### 2.5.2 “Consistency Score” matrix

Next, we evaluated the consistency of the effect of the adaptive classifier on PA for each UC-pair across the 75 datasets. To assess this, we transformed each modulated-PA matrix into a binary matrix, by assigning a score of +1 to positive modulated PA values and 0 to negative or null values. We then summed the binary matrices across all datasets to obtain a consistency score matrix, in which higher values indicate the number of datasets showing improved performance with the adaptive classifier as compared to the static one.

#### 2.5.3 “Modulated Bias” matrix

Non-stationarity affecting EEG signals often leads to the classifier being biased, that is, the erroneous tendency of the classifier to make predictions consistently towards one particular class. By constantly updating its parameters based on incoming signals, adaptive classifiers have been proven effective in reducing the classifier’s bias *(36,62)*.

We evaluated which UC-pair would lead to the lower bias as follows: for each dataset, we considered the bias of the static classifier (static Bias) and the bias of the adaptive classifier (adaptive Bias) for each UC-pair as the absolute value of the difference in the PA between the two classes (Equation (12) - (13)). Then, for each of the 75 EEG datasets, the individual “Modulated bias” matrix was computed as the difference between the adaptive Bias and the static Bias. Last, we calculated the average of all the “Modulated Bias” matrices, yielding to one average “Modulated Bias” matrix across all 75 datasets.

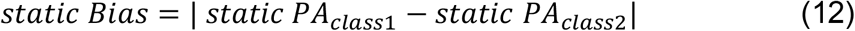

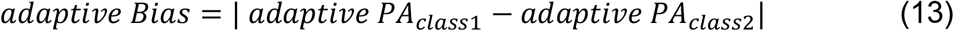

#### 2.5.4 Choice of the optimal UC-pair

Based on these three matrices, we identified one UC-pair to be integrated in the adaptive classifier as the one that maximized the *Modulated PA* and the *Consistency Score* while minimizing the *Modulated Bias*. To quantitatively evaluate the validity of this choice, we conducted a two-tailed paired *t*-test to compare the static PA and the adaptive PA obtained from the pre-recorded datasets by using the select UC-pair. We thus considered for each participant the average PA over the five days of training.

### 2.6 Real-time BCI experiment to evaluate the adaptive classifier’s effectiveness

We then used Study 2 to address our main hypothesis that the improvement obtained with the adaptive LDA classifier by running simulation analysis should translate into better BCI-control in real-time. In this experiment, we employed the same UC-pair as previously identified at the group level based on the Study 1 dataset. We divided our pool of participants into two groups based on the session they performed first leading to an “Adaptive First” group and a “Static First” group.

The BCI experimental protocol was the same as described in the section “BCI loop” for both the static and adaptive classifier sessions, except that during the adaptive session, the classifier in the *online control* phase was updated continuously as described in section 2.4. Every time a participant completed a new trial for each of the two classes (one per syllable) the classifier’s parameters *w* and *b* were re-calibrated using Equations (1) through (7), employing the optimal UC-pair selected from the simulation analysis. This updated classifier was iteratively applied to the forthcoming data.

#### 2.6.1 BCI-control performance

To evaluate performance during the real-time *online control* phase, we computed *BCI-control performance* as the percentage of time where the classifier’s output matched the cued syllables, for each trial independently. Unlike the PA described in the previous sections where each trial is labeled as correctly or incorrectly classified (hit/miss), the *BCI-control performance* is here calculated based on sliding windows of 500 ms and therefore can be considered a more fine-grained way of assessing performance. In addition, as the visual feedback is provided from the classifier’s output, the performance also reflects the subjective perception of control from the individual operating the BCI.

Finally, the overall BCI-control performance for each *online control* phase was determined by averaging values from all trials.

Differences in BCI-control level achieved with the adaptive and with the static classifier were tested with two-tailed paired *t*-tests. To confirm there was no learning between the two sessions, we compared performances during the first and second sessions, irrespective of the order of the classifier used. Last, we compared the performance obtained with the adaptive and the static classifier separately for the “Adaptive First” and “Static First” groups.

Last, we investigated whether the improvement obtained with the adaptive classifier vs. the static one (irrespective of which classifier was used first) could be related to initial BCI performance. More specifically, we hypothesize that those participants who benefited the most from the adaptive classifier were those who exhibited better BCI-control performance during the first session. To test this, we considered the difference in BCI-control performance between the adaptive and static classifiers and computed Pearson’s correlation coefficient with the BCI-control performance observed in the first session.

#### 2.6.2 Cross-Validation Accuracy

To evaluate differences in decoding accuracies between the *offline training* and *online control*, we computed a static LDA classifier separately for the offline and online datasets from both Study 2 experimental recordings (adaptive and static classifier) and tested the accuracy with an 8-fold cross-validation (CV). We checked that there was no difference in the CV accuracy between the two sessions during the offline training as the experimental conditions were the same. We then tested differences in CV accuracy between the *offline training* and *online control*, as a previous study using the same paradigm highlighted a higher decoding accuracy when visual feedback was provided *(33)*. Last, we investigated whether the magnitude of this difference depended on the classifier used (static vs. adaptive). Each of these analyses assessing differences in CV accuracies employed two-tailed paired *t*-tests.

#### 2.6.3 Post-hoc simulation analysis and Individual UC-pair optimization

To assess the reliability of our approach in identifying the optimal UC-values, we performed the same simulation analysis as used for the pre-recorded Study 1 dataset (section 2.5) on the Study 2 dataset. In particular, we evaluated the overlap in the three distinct PA metrics (namely the Modulated PA, the Consistency Score, and the Modulated Bias) used to choose the optimal UC-pair between the two datasets.

Using the online BCI dataset, we further tested the difference in PA obtained for the static vs. adaptive classifier using the selected UC-pair as described in section 2.5 (two-tailed paired *t*-tests).

Previous studies have pointed out that the speed of adaptation is a crucial element for the adaptive classifier as a classifier updating too often might lead to instability in the visual feedback, whereas a classifier updating too slowly might fail to effectively track neural changes due to learning *(63)*, both causing the co-adaptive system to underperform *(64)*. To further optimize the UC parameters, an alternative approach consists in tuning the UC-pair for each individual separately, rather than applying the values computed at group level. Such an approach would, however, require a considerable amount of data and would be computationally time-consuming. We therefore decided not to explore this option in real-time but we evaluated its validity for future developments through a simulation analysis.

We used the same simulation analysis method as detailed in section 2.5, but this time we based the selection of the UC-pair exclusively on the “*Modulated PA*” matrix considering, for each participant, data from both sessions. For each participant, we selected the UC-pair that led to the strongest increase with the adaptive classifier (i.e. the highest value in the “Modulated PA”) and tested for differences with PAs obtained with the static classifier and adaptive with UC-pair obtained at the group level (one-way repeated measures ANOVA). Post-hoc comparisons were carried on with a two-tailed paired *t*-test.

## 3. Results

### 3.1 Simulation analyses to optimize the adaptive LDA classifier

First, we identified the optimal UC_∑_ and UC_μ_ values (i.e. a UC-pair) to be used during real-time BCI-control by performing simulation analysis on a previously recorded dataset employing the same experimental paradigm as the one tested in the present study *(33)* We based the choice of the UC-pair on three distinct metrics: “*Modulated PA*”, “*Consistency Score*”, and “*Modulated Bias*”. The “*Modulated PA*” matrix, obtained as the difference in PA between static and adaptive classification, indicated that combinations yielding the strongest increase in PA (amounting to 5%) with the adaptive classifier fall within a restricted UC range (UC_μ_: 0.4*2^-3^ to 0.4*2^-4^, UC_∑_: 0.4*2^-3^ to 0.4*2^-1^, Figure 3a). Next, we considered the consistency of the adaptive classifier to improve PA and found that the above-mentioned UC range based on the “*Modulated PA*” had the highest consistency score, with 54 out 75 datasets showing an improvement (UC_μ_: 0.4*2^-6^, UC_∑_: 0.4*2^-3^ to 0.4*2^-1^, Figure 3b). Based on both the “*Modulated PA*” & “*Consistency Score*” we selected as the optimal UC parameters for the adaptive LDA classifier to be tested online, a UC_μ_ equal to 0.4*2^-6^ and a UC_∑_ at 0.4*2^-3^ (red dots in Figure 3a-b), and then verified that the chosen parameters were also effective in decreasing the bias between the two classes. Accordingly, the optimized values used with the adaptive classifier allowed a bias reduction of 12.5% as compared to the static classifier (“*Modulated Bias*”, Figure 3c).

**Figure 3:**
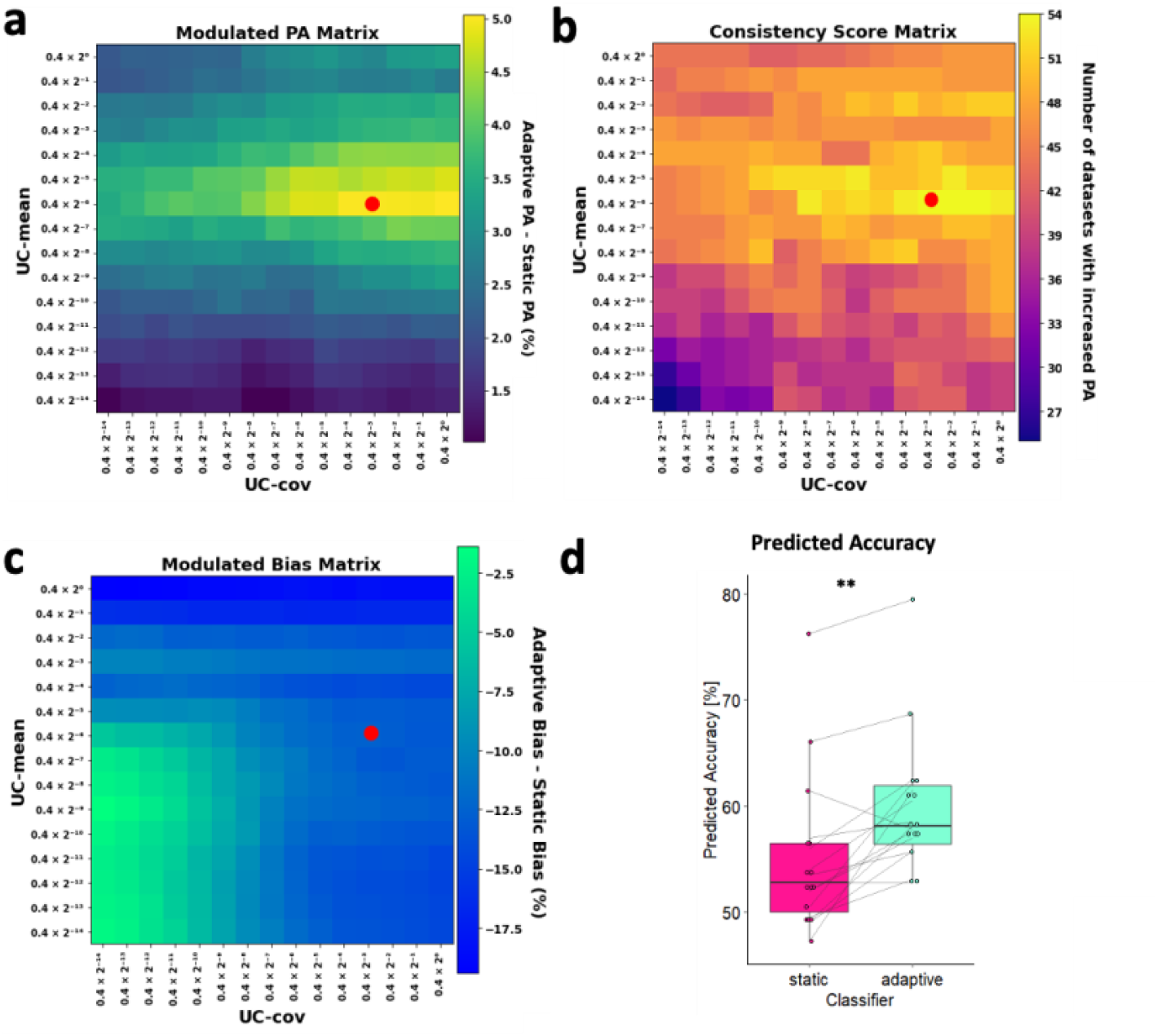
Simulation analyses to optimize the parameters for the adaptive LDA classifier. **(a)-(c):** Results from three metrics considered to evaluate the predicted accuracy using different UC-pair values (UC_∑_ and UC_μ_) using a pre-recorded dataset (Study 1). The x and y axes depict the candidate values range respectively for UC_∑_ and UC_μ_, spanning from 0.4 * 2^-14^ to 0.4, comprising 15 distinct values. **(a)** “Modulated PA” matrix: values (percentage) in this matrix represent the difference between the adaptive PA and the static PA for each UC-pair combination, averaged across datasets. The red dot indicates the maximum value of PA improvement equal to 5% and obtained with UC_μ_= 0.4*2^-6^, and UC_∑_=0.4*2^-3^. **(b)** “Consistency Score” matrix: each element of the matrix indicates the number of datasets where the adaptive PA is higher than the static PA for a given UC-pair. The red dot in correspondence with the UC-pair chosen based on the Modulated PA highlights a value of 54, denoting that these particular UC-parameters yielded improvements in 54 of the 75 datasets. **(c)** “Modulated Bias” matrix: each element of the matrix (percentage) is obtained as the difference in PA between the two classes (i.e. the two syllables) associated with each UC parameter combination. The highlighted value of -12.5% indicates that the PA disparity between the two classes was reduced by 12.5% compared to the static classifier. **(d)** Comparison of the average PA across the 5 datasets (in %) between static and adaptive classifiers with the selected UC-pair (UC_μ_= 0.4*2^-6^, and UC_∑_=0.4*2^-3^, indicated by red dot). The PA obtained was significantly higher for the adaptive (cyan, 60.1 % ± 1.74%) than for the static classifier (pink, 55.1% ± 1.98%). Gray lines connect data from the same participants. Significance is denoted with ** for p < 0.01.

We then quantified whether the decoding improvement due to the adaptive LDA classifier was statistically significant across participants and found that this was the case (T_14_= 4.29, p < 0.01, d = 1.1, Figure 3d). Out of 15 participants, 13 exhibited a higher PA with the adaptive LDA classifier than with the static classifier.

### 3.2 Real-time BCI experiment

#### 3.2.1 BCI-control performance

We tested the effectiveness of using the selected UC-pair in a real-time setting with a new group of participants, who performed two separate BCI-control sessions, one employing a static LDA and the other an adaptive LDA classifier, in randomized order. Two-tailed paired *t*-tests revealed higher BCI-control performance with the adaptive than the static classifier (T_19_=2.99, p<0.01, d= 0.67, Figure 4a). This effect was present in 15 out of 20 participants and did not depend on the order in which each of the classifiers was used (“Adaptive First” and “Static First” groups). Both groups had a statistically significant improvement with the adaptive classifier (“Adaptive First” Group: 54.3 % ± 1.56 % for the adaptive classifier, 52.3 % ± 2.11 % for the static classifier, T_9_= 2.27, p<0.05, d= 0.72; “Static First” Group: 57.2 % ± 2.31 % with the adaptive classifier, 52.3 % ± 0.35 % for the static classifier, T_9_=2.34, p<0.05, d= 0.74).

**Figure 4:**
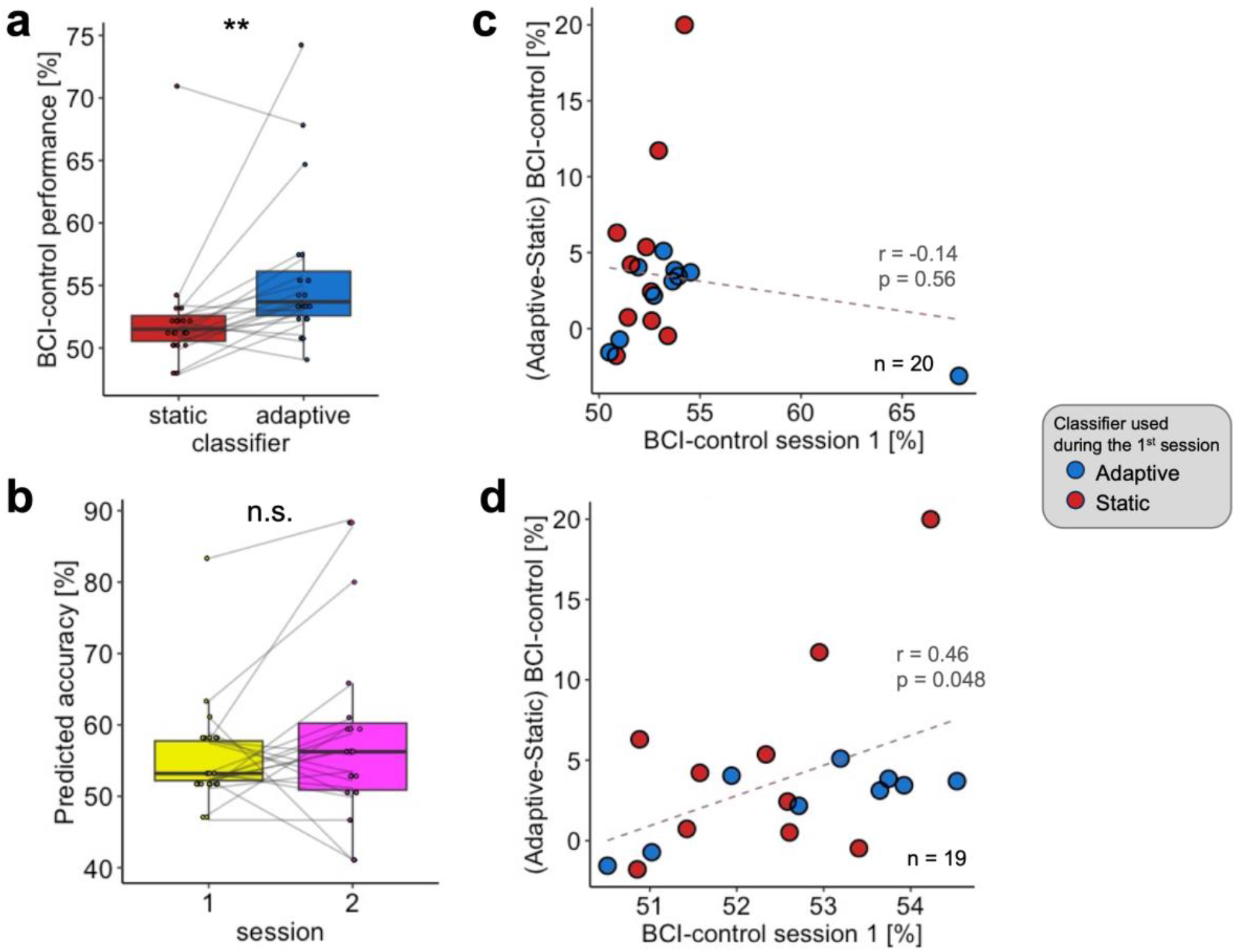
Impact of the adaptive LDA classifier on BCI-control performance. **(a)** The adaptive LDA classifier led to better BCI-control performance (55.7% ± 1.39%) relative to the static classifier’s (52.3% ± 1.04%). This effect was present in 15 out of 20 participants. **(b)** There was no significant difference in BCI-control performance between the first (53.3% ± 0.81%) and the second session (54.7% ± 1.62%), irrespective of the classifier used. Individual dots represent individual BCI-control performance, gray lines link data from the same participant. **(c)-(d)** Correlation between BCI-control performance as observed during the first session (x-axis) and difference in performance between adaptive and static classifiers (y-axis). The gray dotted lines illustrate the regression line. Results are shown considering data from all participants **(c)** and with one outlier removed **(d)**. Significance is denoted with ** for p < 0.01.

Furthermore, we also controlled for learning across sessions and found no difference in BCI-control between the first (53.3 % ± 0.81 %) and second session (54.7 % ± 1.62 %, T19 = 1.07, p = 0.3, d = 0.24, Figure 4b).

To test whether participants with better initial performance benefited more from the adaptive classifier than the poorer performers, we correlated.

We correlated the difference in BCI-performance between the adaptive and static classifier with the BCI-performance of the first session and found no statistically significant relationship (Figure 4c, ρ = -0.138, p = 0.56). However, this statistical result was driven by one outlier point, who had the highest performance of all recordings with the static LDA classifier (static: 70.9 %, adaptive: 67.8 %). Upon removing this value, the correlation coefficient reached statistical significance (Figure 4d, ρ = 0.458, p = 0.048).

#### 3.2.2 Cross-validation accuracy

As expected, we found no difference in the cross-validation (CV) accuracy obtained by classifying data from the *offline training* phase between the adaptive (49.7% ± 0.72%) and the static classifier (50.4% ± 0.88%) (T_19_=0.59, p=0.56, d=0.13, Figure 5a). Surprisingly, there was no significant difference in the CV accuracy for the *online control* phase (Figure 5b, T_19_=1.4, p=0.18, d=0.31). We then tested for a decoding improvement from the *offline training* to the *online control*, separately for the adaptive and static sessions, and found an increase for both sessions (Static classifier: Offline CV = 50.4% ± 0.88%, Online CV = 53.9% ± 1.7%, T_19_=2.95,p<0.01, d=0.66; Adaptive classifier: Offline CV = 49.7% ± 0.72%, Online CV = 56% ± 1.95%, T_19_=3.11,p<0.01, d=0.7, Figure 5c-d). The critical test was whether the improvement in the online setting was potentiated by the adaptive classifier, i.e. whether there was an added value of an adaptive classifier in a closed loop setting. We found a trend for a stronger increase associated with the use of the adaptive classifier (adaptive: 6.3% ± 2%, static: 3.5% ± 1.18%, T_19_=1.85, p=0.08, d=0.41, Figure 5e).

**Figure 5:**
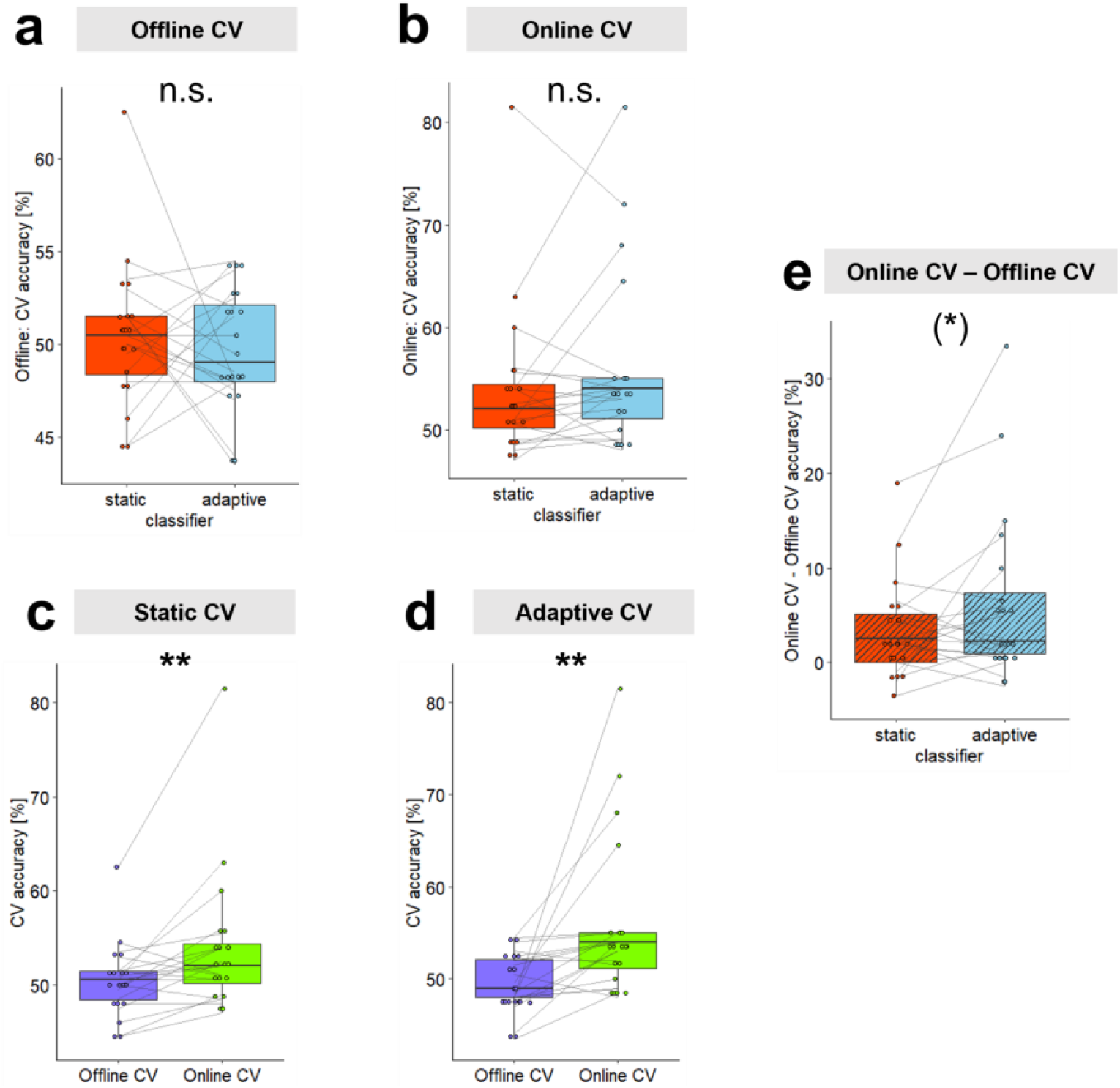
Cross-validation accuracy comparisons. We considered the cross-validation accuracy of decoding models obtained from a static classifier in order to compare decoding from different datasets (offline vs online, static vs adaptive). **(a)** There was no significant difference in the CV accuracy obtained from the *offline training* phase between the static (orange, 50.4% ± 0.88%) and the adaptive classifier (light blue, 49.7% ± 0.72%) sessions, **(b)** nor for the *online control* phase (static: orange, 53.9% ± 1.7%; adaptive classifier, light blue: 56% ± 1.95%). **(c)-(d)** Data acquired during the *online control* phase (i.e. when the user received a real-time feedback, green) led to significantly higher CV accuracy than during the *offline training* phase for both the static classifier session (**c**, online: 53.9% ± 1.7%, offline CV: 50.4% ± 0.88%) and the adaptive classifier session (**d**, online: 56% ± 1.95%, offline: 49.7% ± 0.72%). **(e)** We tested for differences between the static and adaptive BCI session in the amount of CV accuracy improvement from the *offline training* to the *online control* phase and found a trend toward significance (adaptive: 6.3% ± 2%, static: 3.5% ± 1.18%). Dots represent individual participants’ data. Gray lines link data from the same participant. Significance is denoted with ** for p < 0.01, and (*) for p=0.08.

#### 3.2.3 Post-hoc simulation analysis

We then investigated whether the selected UC-pair used for the adaptive LDA classifier and derived from the pre-recorded dataset aligned with that obtained from the real-time experiment. We applied the same simulation analyses as described in section 2.5. to compute the “*Modulated PA*”, “*Consistency Score*” and “*Modulated Bias*” for this new dataset. The selected UC-pair felt within optimal PA regions of the real-time experiment. More specifically, the selected UC-pair was the second pair in terms of increased accuracy with the adaptive classifier as compared to the static classifier (“*Modulated PA*”, Figure 6a). Such an improvement using the selected PA appeared in 26 out of 40 datasets, as indicated in the “*Consistency Score*” matrix, close to the maximum value of 29 datasets (Figure 6b, black dot). Of note, the UC-pair that led to the highest consistency value was associated with a lower improvement in PA than the UC-pair used in the real-time experiment (red dot in the “*Modulated PA*” matrix). The selected UC-pair also reduced the bias between the two classes of -10.92 %, the second strongest reduction throughout the entire “*Modulated Bias*” matrix (Figure 6c).

**Figure 6:**
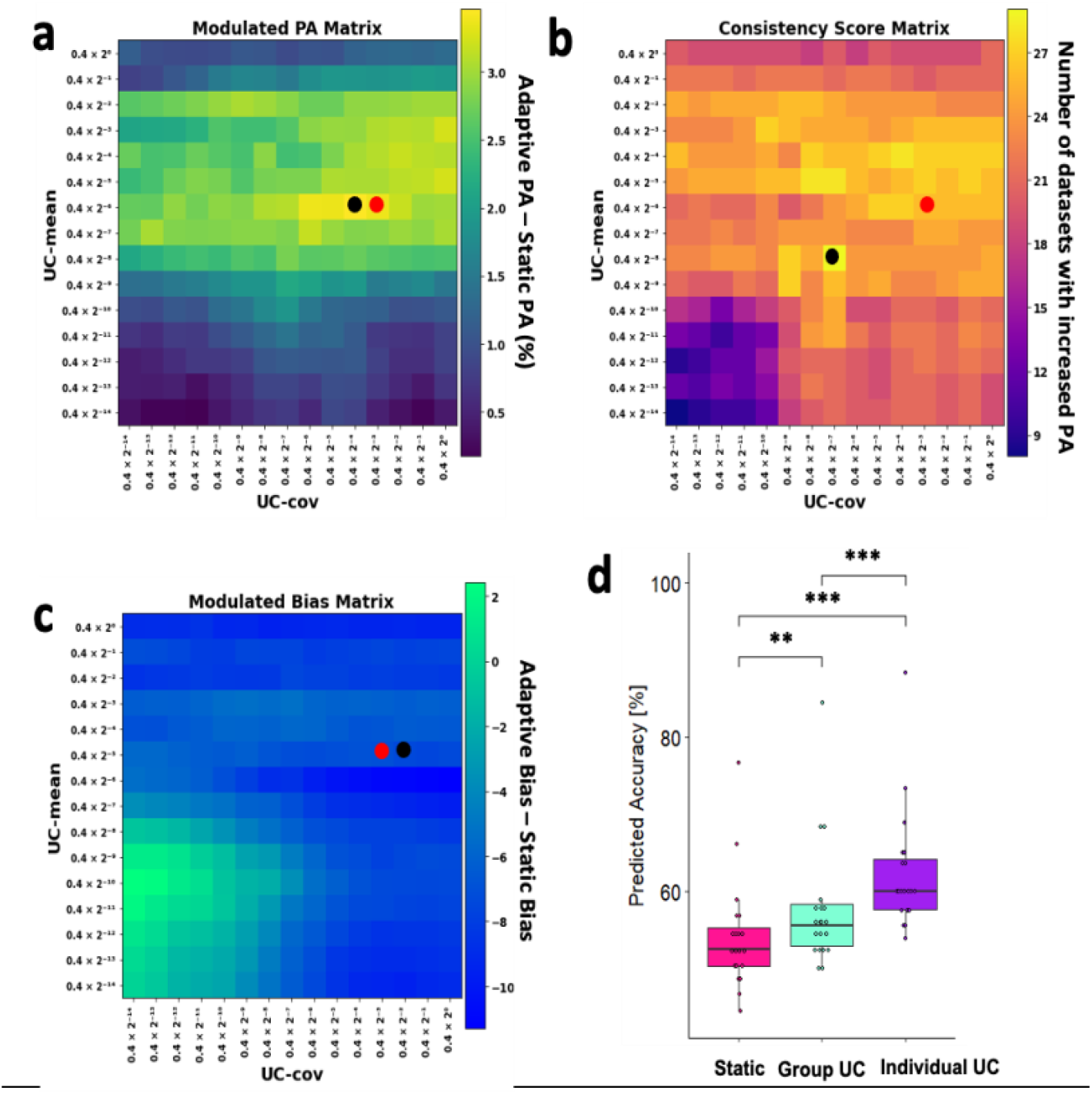
Simulation analyses on the real-time experiment dataset. **(a) - (c)** Predicted accuracy for different UC values computed with classification simulation on the real-time experiment dataset (Study 2). The x and y axes indicate the candidate range for UC_∑_ and UC_μ_ respectively, spanning from 0.4 * 2^-14^ to 0.4, comprising 15 distinct values. Red dots represent the selected UC parameters used in the real-time experiment (selected UC-pair: UC_μ_= 0.4*2^-6^, UC_∑_=0.4*2^-3^). Black dots represent the local maximum/minimum value in each matrix. **(a)** “*Modulated PA*” matrix: by using the selected UC-pair (red dot), the PA increased by 3.36% compared to the static classifier’s PA. The maximum increase was 3.45% (black dot). **(b)** “*Consistency Score*” matrix: the selected UC-pair (red dot) led to a PA improvement in 26 out of the 40 datasets. The maximum value in this matrix was 29 (black dot). **(c)** “*Modulated Bias*” matrix: the selected UC-pair was effective in decreasing the bias between the two classes by 10.92%, and the maximum bias reduction was -11.14% (black dot). **(d)** Comparison of simulated analysis performance across different classifiers. Each point marks the PA of an individual participant. A one-way repeated measures ANOVA confirmed a significant main effect of classifier type on the PA. Subsequent post-hoc paired *t*-tests showed that the PA for the adaptive classifier with individual UC (62.2% ± 1.73%, purple) was higher than both the static classifier (54.1% ± 1.59%, pink) and the adaptive classifier with Group UC (57.5% ± 1.8%, green). Performance with the adaptive classifier with group UC exceeded the PA obtained with the static classifier. Significance is denoted with ** for p < 0.01, *** for p < 0.001.

Two-tailed paired *t*-tests showed an improvement in PA using the adaptive classifier with selected UC-pair relative to the static classifier, consistently with the results from the same simulation analysis conducted on the previously acquired Study 1 dataset (Figure 6d, T_19_=3.2, p<0.01, d=0.72).

### 3.3 Individual UC-pair optimization

Finally, we investigated whether adapting the UC-pair for each participant further boosted PA. A one-way repeated measures ANOVA with classifier type as a factor (static, adaptive with the UC-pair optimized at the group level, and adaptive with the UC-pair optimized at the individual level) showed a significant main effect (F_(2,38)_ = 43.8, p < 0.001, η^2^_p_ = 0.166). Post-hoc paired *t*-tests with Bonferroni corrections revealed that the PA obtained with the individual UC-pair was statistically significantly higher than the static classifier (T_19_=8.72, Bonferroni-adjusted p < 0.001) and the group UC-pair (T_19_ = 8.65, Bonferroni-adjusted p < 0.001) (Figure 6d).

## 4. Discussion

In this study, we explored whether employing an adaptive LDA classifier for decoding the imagery of two syllables from EEG signals would yield improved BCI-control compared to a static classifier, commonly used in the field. To do this, we calibrated the adaptive LDA based on simulation analyses performed on a pre-recorded dataset, and then tested its effectiveness in a new experiment in which healthy volunteers performed two sessions of BCI-control, one employing a static LDA and the other an adaptive LDA classifier, over two separate days.

As hypothesized, we found that the adaptive classifier enhanced BCI-control performance in the real-time BCI experiment. This effect was not related to participants learning to operate the BCI system, as no difference was found between the first and the second experimental session, and it was present irrespective of the classifier used during the first session. These findings support the benefit of using adaptive classifiers in the field of imagined speech decoding, which is notoriously characterized by low online decoding accuracy, hardly exceeding the chance level (23,33,65).

Furthermore, we found no difference in decoding accuracy computed in post-processing (i.e. CV accuracy) between the static and the adaptive session, neither for the offline training, nor for online control datasets. This result was expected for the offline training (Figure 5a), the first part of the experiment preceding the classifier training when no feedback is provided to the user (thus characterized by the same experimental conditions). However, it was surprising to observe no difference for the online training dataset (Figure 5b): since we measured higher BCI-control performance when computed in real-time using the adaptive classifier, it would have been plausible to find such a difference also when decoding was applied in post-processing using a different approach. This discrepancy suggests that the improvement observed in real-time is solely due to an optimization on the decoder side and it is not accompanied by adaptation of the underlying neural activity when using one or the other classifier. If a difference had emerged along the course of the experiment, it would have been likely due to the neural data being more discriminable (between the two classes) during the adaptive session. Ultimately, this would be reflected in a higher CV accuracy.

Confirming previous results using the same paradigm (33), we found higher decoding accuracy for signals recorded during the online control than the offline training, indicating that the real-time feedback helped users to self-regulate brain patterns. Interestingly, the extent of decoding improvement when the imagery task is performed within a closed loop (online control) tended to be stronger in the adaptive session than in the static one. This suggests that, if not over the course of a single online control session, boosting real-time control with an adaptive classifier has the potential of enhancing neural discriminability in the medium term. Enhanced benefits from the real-time feedback with an adaptive classifier might also accelerate learning to operate the BCI. The acquisition of BCI-control skills has been shown to be possible using a static classifier by training participants over 5 consecutive days (33). We can thus reasonably expect that an adaptive classifier further promotes learning by capitalizing on the co-adaptation between the decoders and the user, previously shown to induce neural changes on a short time scale (66).

Enhancing the accuracy of current decoders could further boost metacognitive abilities that play a key role in learning to consciously self-regulate ongoing brain-activity (67,68). This conscious aspect, together with an unconscious learning, originally posited by Lacroix as the dual-process model (69), has been proposed as a theoretical framework to account for learning BCI control (68). Similarly, improving decoding will impact higher-order mechanisms related to action execution such as the subjective sense of being in control of BCI-actions: this fundamental mechanism has been characterized for motor imagery BCI-control as relying mainly on the congruency between intentions and sensory feedback (70,71). By enhancing these higher-order cognitive mechanisms, a more accurate decoding from the machine side could further boost BCI-control on the user side in a synergistic fashion.

Our experimental results also show that the approach we used to identify the optimal update parameters at the group level is robust and reliable, as we found matching values using the pre-recorded dataset and the real-time dataset acquired specifically to validate our method. Albeit effective, we found that the UC values chosen at the group level did not benefit all participants, as 5 out of 20 did not improve BCI-control with the adaptive classifier. We then explored whether optimizing the UC values at the individual level resulted in a more uniform increase and found that individualizing the UC parameters improved PA in all participants. Although promising, individualized simulations require substantial amounts of data and computational resources, thus their effectiveness remains to be established in real-time control settings.

Furthermore, the fact that the adaptive classifier tested in real-time was not optimally tailored to each participant’s individual characteristics might explain the absence of a clear link between initial BCI-control abilities and the improvement associated with the use of an adaptive classifier that we initially hypothesized. In fact, we found such a positive trend to be present but heavily affected by one outlier who performed surprisingly well with the static classifier. Further investigations are needed to identify inter-individual differences in the ability to exploit the full potential of an adaptive classifier.

Together with the individual tailoring of the update speed (i.e. the UC value), another potential optimization of the adaptive classifier consists in varying UC-values in the course of the experiment, instead of being kept constant as in our study. Previous studies have put forward the use of Kalman adaptive classifier, whereby the Kalman gain would vary the UC values according to the classifier’s output in real-time in a supervised fashion *(36,37)*. More specifically, if the classifier output is accurate, the system will rely more on the current decoding model, slowing the update speed.

Conversely, if the classifier predictions are inaccurate, the update speed will be increased *(49,50,72)*. This approach could constitute a further improvement of the decoder, together with integrating a feature selection step that would focus the decoding on specific regions (left central and temporal bilateral) and frequency bands (alpha to low-gamma), shown to be the most discriminant with surface EEG for syllables imagery *(33)*.

## 5. Conclusions

The present study shows for the first time the effectiveness of using an adaptive classification approach to improve BCI-control based on imagined speech from surface EEG, notoriously known to be affected by low performance. Such technical efforts might appear marginal with regards to recent advances in decoding attempted speech from intracranial implants with rates approaching half the normal speech *(12,14)*. While non-invasive systems will presumably never constitute a means of communication, simple yet effective binary BCIs can in fact be instrumental in enabling users to learn self-regulating brain patterns in preparation for operating more complex BCI systems. This has been shown in the context of an intermediate phase that forms a part of progressive training, viewed from the standpoint of human learning (73). Using such a progressive training method, previous studies achieved high BCI performances in motor imagery tasks (74–77) and the learning previously observed (33) suggest this will hold true for imagined speech. Moreover, the use of adaptive classifiers is particularly relevant for surface-EEG recordings as an effective means to increase the number of decoding classes. Most of the current BCIs are limited to binary classification due to the necessity of having enough training data to counterbalance the low signal-to-noise ratio. By using data from the real-time control as training data, an adaptive classifier shortens the training time and favors the development of multi-class BCIs (78,79). In summary, our results encourage the systematic use of adaptive decoders in the field of speech-BCI, a compelling, robust and easy-use technological solution that presents numerous advantages without any additional computational costs.

## Author Contributions

Conceptualization, Silvia Marchesotti, Anne-Lise Giraud, Shizhe Wu, ; methodology, Shizhe Wu.; software, Shizhe Wu, Kinkini Bhadra.; formal analysis, Shizhe Wu, Silvia Marchesotti.; investigation, Shizhe Wu, Kinkini Bhadra.; writing—original draft preparation, Shizhe Wu, Silvia Marchesotti, Anne-Lise Giraud; writing—review and editing, Silvia Marchesotti, Anne-Lise Giraud, Shizhe Wu, Kinkini Bhadra; visualization, Shizhe Wu, Silvia Marchesotti.; supervision, Silvia Marchesotti, Anne-Lise Giraud; project administration, Silvia Marchesotti, Anne-Lise Giraud.; funding acquisition, Anne-Lise Giraud. All authors have read and agreed to the published version of the manuscript.

## Institutional Review Board Statement

The study was conducted in accordance with the Declaration of Helsinki, and approved by the local Ethics Committee (Commission cantonale d’éthique de la recherche, project 2022-00451).

## Informed Consent Statement

Informed consent was obtained from all subjects involved in the study.

## Data Availability Statement

The data that support the findings of this study are available from the corresponding author upon reasonable request.

## Acknowledgments

We thank the Human Neuroscience Platform of the Fondation Campus Biotech for technical advice.

